# Dynamic phosphorylation of Hcm1 promotes fitness in chronic stress

**DOI:** 10.1101/2024.09.18.613713

**Authors:** Michelle M. Conti, Jillian P. Bail, Aurelia R. Reynolds, Linnea G. Budge, Mackenzie J. Flynn, Rui Li, Lihua Julie Zhu, Jennifer A. Benanti

## Abstract

Cell survival depends upon the ability to adapt to changing environments. Environmental stressors trigger an acute stress response program that rewires cell physiology, downregulates proliferation genes and pauses the cell cycle until the cell adapts. After the acute response is resolved, cells resume cycling but at a reduced rate. The importance of cell cycle changes for survival in chronic stress is not clear. Here, we show that dynamic phosphorylation of the yeast cell cycle-regulatory transcription factor Hcm1 is required to maintain fitness in chronic stress. Hcm1 is activated by cyclin dependent kinase (CDK) during S-phase and is inactivated by the phosphatase calcineurin (CN) in response to stressors that signal through increases in cytosolic Ca^2+^. Expression of a constitutively active, phosphomimetic Hcm1 mutant reduces fitness in stress, suggesting Hcm1 inactivation is required for survival. However, a comprehensive analysis of Hcm1 phosphomutants revealed that Hcm1 activity is also important to survive stress, and that all mutants with fixed phosphorylation states are less fit in stress. Moreover, pulses of Hcm1 activity are necessary to maximize target gene expression in stress. These findings demonstrate that the expression levels of Hcm1 target genes influence fitness in stress and suggest that the dynamic phosphorylation of cell cycle regulators plays a crucial role in promoting survival in stressful environments.

## Introduction

Cells are continuously exposed to stressors in the environment and must adapt to these challenges to survive and proliferate. Adaptation not only protects healthy cells from death, but conversely, it can promote the development of disease. For instance, cancer cells must adapt to stressful environments when they metastasize to distant sites [1,2], and fungal pathogens rely upon stress response pathways for survival within the host [3]. Despite the importance of this process, the long-term changes that cells must undergo to maintain fitness and proliferation when faced with chronic stress are poorly understood.

The acute stress response, which occurs immediately following exposure to an environmental stressor, is conserved from yeast to humans and includes a downregulation of protein synthesis and an upregulation of stress response genes [4,5]. In addition to these changes that impact cell physiology, cell cycle-regulatory genes are downregulated, and cells undergo a transient cell cycle arrest [6,7]. After cells adapt to the new environment, the acute stress response is resolved and cells resume proliferation in the new environment, albeit at a reduced rate [8–10]. Transient cell cycle arrest during the acute stress response is thought to be crucial to promote long-term survival [9–11]. However, the importance of cell cycle changes for survival in chronic stress is not well understood.

Some stressors signal through an increase in cytosolic Ca^2+^ and coordinate the stress response program and cell cycle changes by activating the conserved Ca^2+^-activated phosphatase calcineurin (CN) [12,13]. CN activation leads to a decrease in expression of cell cycle-regulatory genes and controls the length of cell cycle arrest, in combination with the stress-activated MAPK Hog1/p38 [14]. One direct target of CN in budding yeast is the S-phase transcription factor (TF) Hcm1 [15]. Hcm1 is a forkhead family transcriptional activator which, like it’s human homolog FoxM1, plays a crucial role in maintaining genome stability [16,17]. Hcm1 controls expression of key cell cycle genes including histone genes, downstream cell cycle-regulatory TFs, and genes that regulate mitotic spindle function. As cells progress through the cell cycle, Hcm1 is activated by multisite phosphorylation by cyclin dependent kinase (CDK) [18]. CDK phosphorylates eight sites in the Hcm1 transactivation domain (TAD) to stimulate its activity and a three site phosphodegron in the N-terminus to trigger proteasomal degradation. Immediately following exposure to CaCl2 or LiCl stress, two stressors that activate CN, CDK activity decreases, the activating phosphates on Hcm1 are removed by CN, and expression of Hcm1 target genes decreases [14,15,19]. Whether or not Hcm1 inactivation is critical for cells to adapt and survive in the face of chronic stress is unknown.

Mutations within the Hcm1 TAD have been used to study the consequences of phosphorylation. Phosphomimetic mutations at all CDK phosphosites in the TAD generate a constitutively active protein that leads to increased expression of Hcm1 target genes [18]. In normal growth conditions this mutant provides a fitness advantage to cells, rendering them more fit than wild type (WT) [15,20]. This is a surprising result because advantageous mutations are expected to be selected for during evolution. However, the fact that activating phosphates are removed by CN when cells are faced with stress suggests that dephosphorylation and inactivation of Hcm1 may be necessary for cells to survive in stressful environments.

Here, we investigated this possibility and found that cells expressing a constitutively active, phosphomimetic Hcm1 mutant lose their fitness advantage when they proliferate in LiCl stress for up to 30 generations. To determine the optimal level of Hcm1 activity for fitness in this environment, we screened a collection of Hcm1 mutants that encompass all possible combinations of non-phosphorylatable and phosphomimetic mutations in the TAD, representing the entire spectrum of possible activity levels. Surprisingly, this screen revealed that almost all mutants were less fit, relative to WT, when growing in stress compared to stress-free conditions. Moreover, Cks1-priming sites that stimulate Hcm1 activity by promoting phosphorylation by CDK became more important for fitness when cells were grown in chronic stress. Finally, mutants that have increased Hcm1 activity because proteasomal degradation is blocked, but retain dynamic phosphoregulation of the TAD, upregulated Hcm1 target gene expression to a greater extent and were more fit than WT cells in chronic stress. These results demonstrate that simple Hcm1 inactivation is not the mechanism by which cells survive in chronic stress; instead, dynamic regulation of the Hcm1 activity – obtained through a combination of phosphorylation by CDK and dephosphorylation by CN – is critical to maintain fitness.

## Results

### Expression of a phosphomimetic Hcm1 mutant decreases fitness in LiCl stress

CDK phosphorylates eight sites in the Hcm1 TAD to activate the protein during an unperturbed cell cycle [18]. When cells are exposed to a CN-activating stressor such as CaCl2 or LiCl, these phosphates are removed by the phosphatase CN [15] (Fig 1A). This dephosphorylation occurs rapidly after exposure to LiCl [15] and, notably, Hcm1 remains in a hypophosphorylated state after cells adapt to LiCl stress and resume cycling (Fig 1B). To determine if dephosphorylation of Hcm1 is necessary for cells to survive when faced with chronic LiCl stress, we utilized a constitutively active phosphomimetic Hcm1 mutant, Hcm1-8E, in which each CDK site in the TAD is mutated to two glutamic acids (S/T-P to E-E) to mimic the charge of a phosphate [18,20]. The Hcm1-8E protein is more active than WT Hcm1 and, as a result, confers a fitness advantage to cells in a competitive growth assay in optimal growth conditions [15,20]. To determine the consequence of elevated Hcm1 activity in stress, we used the same competitive growth assay to compare the fitness of cells expressing Hcm1-8E to cells expressing WT Hcm1, in the presence or absence of LiCl (Fig 1C-1D, S1A-S1C Fig). As previously shown [20], *hcm1-8E* cells exhibited a fitness benefit in the absence of stress (Fig 1C). However, *hcm1-8E* cells lost their fitness advantage and displayed a modest fitness defect when cultured in medium containing LiCl (Fig 1D), supporting the possibility that cells need to inactivate Hcm1 to survive in the presence of chronic LiCl stress.

**Figure 1.**
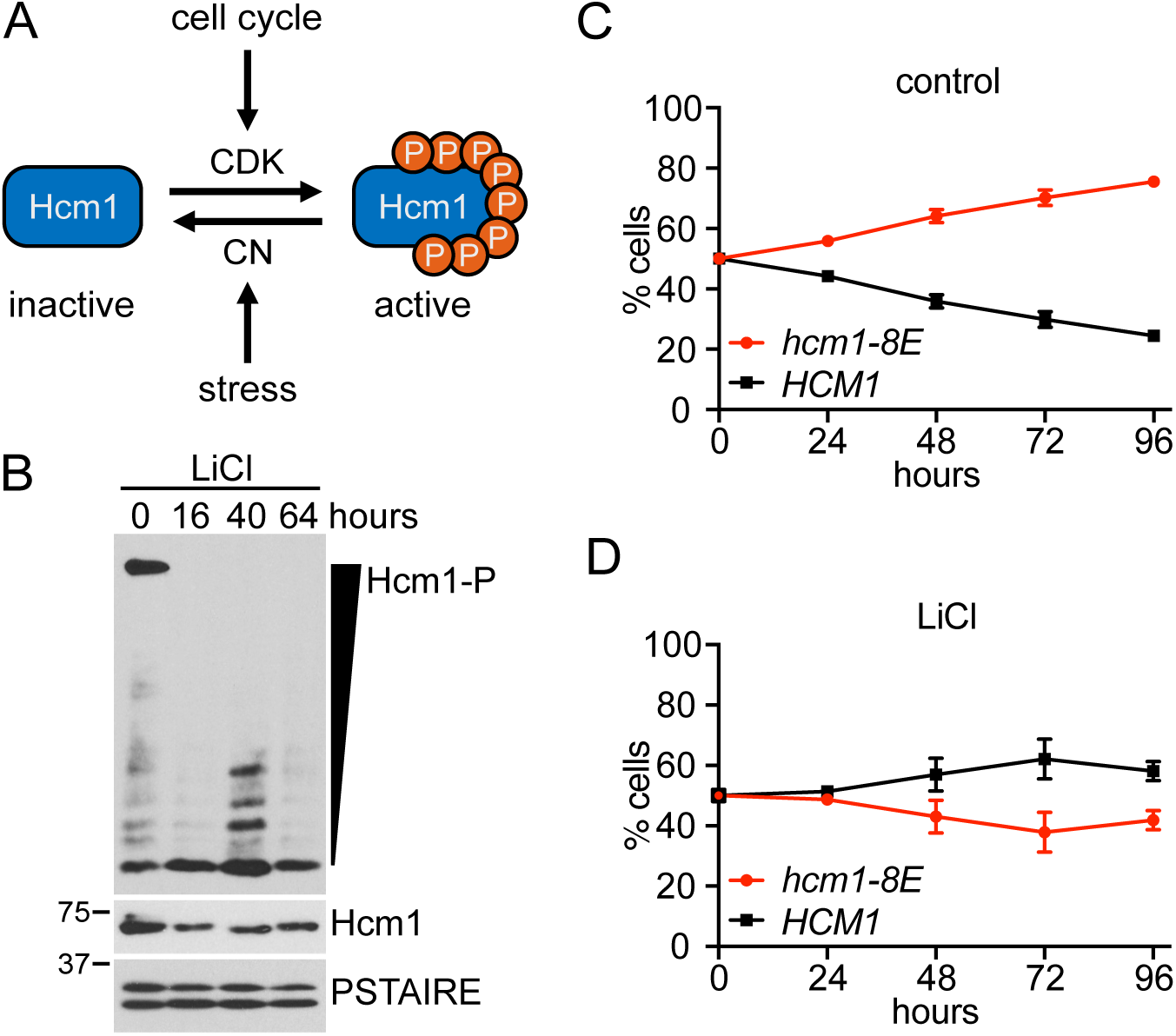
Expression of a phosphomimetic Hcm1 mutant decreases fitness in LiCl stress. (A) Hcm1 activity is regulated by cyclin-dependent kinase (CDK) and the phosphatase calcineurin (CN). (B) Phos-tag and standard Western blots showing Hcm1 phosphorylation and expression after the indicated number of hours in LiCl stress. Hcm1 was detected with an antibody that recognizes a 3V5 tag, PSTAIRE is shown as a loading control. Representative blots from n=3 experiments are shown. (C-D) Strains with the indicated genotypes were co-cultured in control media (C) or media with 150mM LiCl (D). Percentage of each strain was quantified by flow cytometry at the indicated timepoints. An average of n=3 biological replicates is shown. Error bars represent standard deviations.

### Hcm1 mutants with fixed phosphorylation states are less fit in LiCl stress

Hcm1 retains some phosphorylation when cells are grown for three days in LiCl (Fig 1B), suggesting that some activity may remain, and that a reduced level of Hcm1 activity might be optimal in stress. To test this hypothesis, we employed Phosphosite Scanning, a recently developed approach that can simultaneously determine the effects of hundreds of phosphosite mutations on cellular fitness [20]. Phosphosite scanning was previously used to screen a collection of Hcm1 mutants in which each CDK phosphosite (S/T-P) in the TAD is mutated to an unphosphorylatable alanine (A-P) or two glutamic acids (E-E), in all possible combinations (A/E library, Fig 2A) [20]. In unstressed growth conditions, fitness values conferred by these Hcm1 mutants are highly correlated with the activity of each mutant and this collection of mutants represents the entire continuum of possible Hcm1 activities, from the completely inactive Hcm1-8A mutant (with all phosphosites mutated to A-P) to the Hcm1-8E mutant that has increased activity relative to WT [20]. To determine the optimal amount of activity in stress, we used the same Phosphosite Scanning approach and screened the Hcm1 A/E library in control and LiCl containing media in parallel (Fig 2B).

**Figure 2.**
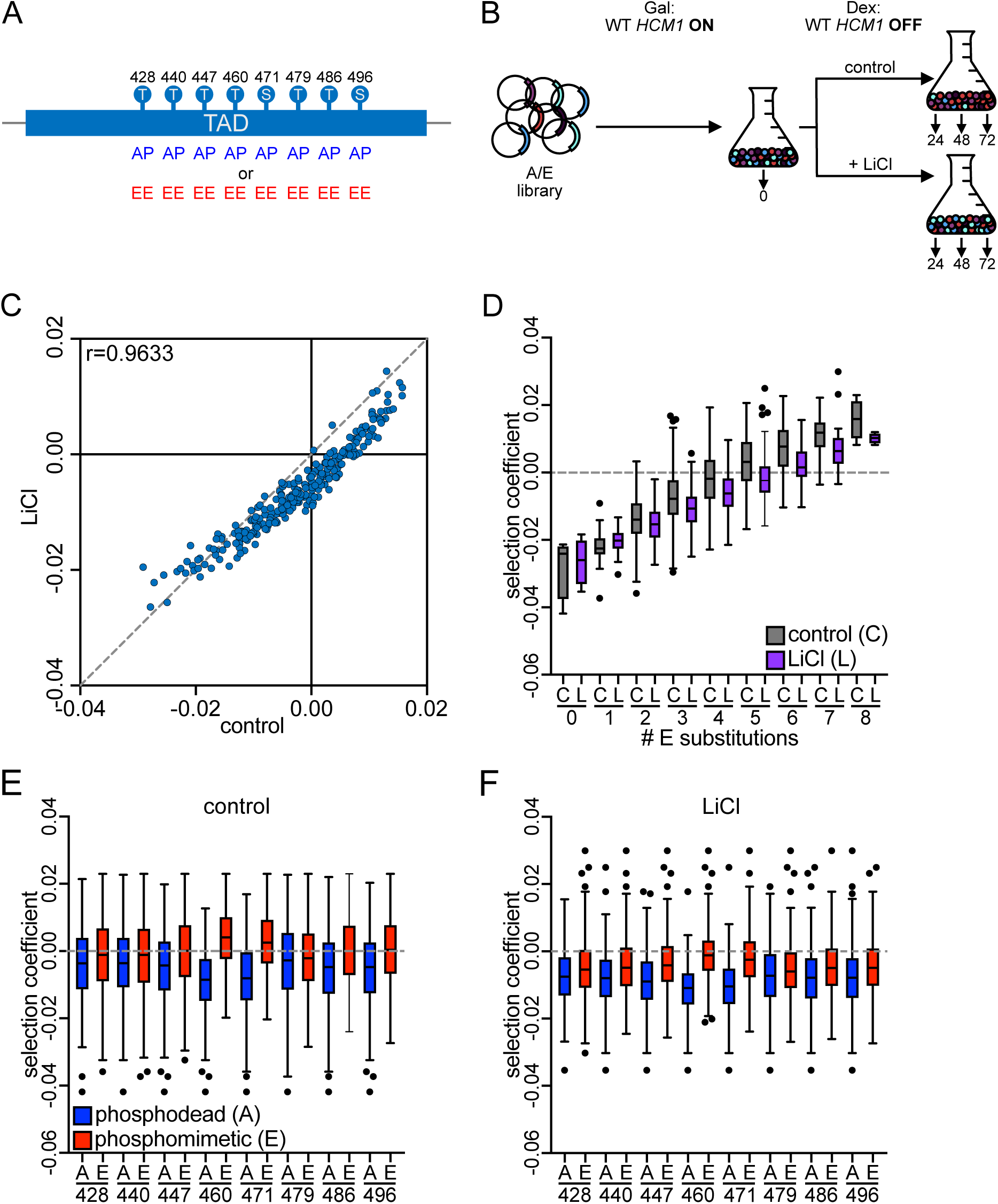
Phosphosite mutations in Hcm1 decrease fitness in stress. (A) Schematic depicting the Hcm1 A/E library. (B) Schematic of Phosphosite Scanning screens. Plasmids expressing all 256 mutants in the A/E library, as well as WT *HCM1*, were transformed into a strain in which expression of the genomic copy of *HCM1* is controlled by a galactose-inducible promoter. The pooled population growing in galactose was diluted and split into dextrose containing media (to shut-off expression of the endogenous copy of WT *HCM1*) with or without 150mM LiCl at the start of the experiment. (C) Scatterplot comparing average selection coefficients for each mutant in the A/E library in control and LiCl media. Pearson correlation (r) is indicated. (D-F) Box and whisker plots comparing the selection coefficients of different groups of mutants. The black center line indicates the median selection coefficient, boxes indicate the 25^th^-75^th^ percentiles, whiskers represent 1.5 interquartile range (IQR) of the 25^th^ and 75^th^ percentile, black circles represent outliers. In all panels, selection coefficients are an average of n=4 biological replicates. (D) shows selection coefficients in cells with the indicated number of phosphomimetic mutations in control or LiCl conditions. (E-F) show selection coefficients of mutants that are either phosphodead or phosphomimetic at each position, in control (E) or LiCl containing medium (F). Statistics for panels D-F are included in S2 Dataset.

As in the previous study, the Hcm1 A/E library was transformed into a strain where the genomic copy of Hcm1 was expressed from a galactose-inducible promoter. At the start of the screen, cells growing in galactose were diluted into control medium or medium containing LiCl and expression of WT Hcm1 was shut off by adding dextrose (S1E Fig), ensuring that all Hcm1 protein in the cells was derived from the plasmid library (Fig 2B). Cultures were periodically sampled and diluted over the course of 72 hours, and the relative abundance of each mutant was tracked via sequencing, as described previously [20]. Unlike the previous study where cells were diluted every 12 hours to maintain logarithmic growth, cells were diluted every 24 hours, which allowed cells in control conditions to approach saturation and exit the cell cycle (S1D Fig). However, Hcm1 expression resumed following each dilution when cells reentered the cell cycle (S1E Fig), and selection coefficients of mutants screened in control conditions using 12- and 24-hour intervals were highly correlated (S2A-C Fig). Thus, the modified screening protocol accurately reflects the relative fitness values of Hcm1 mutants.

We first investigated how overall fitness was impacted when cells were growing in stress. Selection coefficients of each mutant in LiCl-containing medium were directly compared with those from control medium (Fig 2C). Surprisingly, although there was a strong correlation between selection coefficients in the two environments, almost all mutants showed a stronger reduction in fitness, relative to WT, in LiCl conditions than they exhibited in control conditions. This effect was most severe for mutants with greatest number of phosphomimetic (activating) mutations, whereas mutants that had two or fewer phosphomimetic mutations were most similar between control and LiCl media (Fig 2D, S2D Fig). Notably, although the *hcm1-8E* mutant displayed a modest fitness defect in pairwise competition assays carried out in LiCl (Fig 1D), it had a slight fitness advantage in pooled screens (Fig 2D, mutant with eight E substitutions). This observation is consistent with previous findings that pooled screens result in higher selection coefficients than pairwise competition assays and is likely due to technical differences in the experimental approach [20]. However, the *hcm1-8E* mutant fitness advantage was modest in pooled LiCl screens and decreased in LiCl compared to control conditions. These data demonstrate that, contrary to our expectation, Hcm1 activity is required for fitness in stress. However, the most highly active mutants exhibit reductions in fitness compared to control growth conditions.

Next, we wanted to determine how individual TAD phosphosites impact cellular fitness in LiCl stress. To do this, we compared the selection coefficients of all mutants that had either a non-phosphorylatable alanine (A) or two glutamic acids (E) at each site. In both control and LiCl conditions, a phosphomimetic mutation at any site increased fitness relative to an alanine substitution at the same site (Fig 2E-2F, compare red and blue boxes). Notably, mutations at sites T460 and S471 had the greatest effect on fitness in both conditions, consistent with previous measurements in the absence of stress [20]. Together, these data show that when phosphorylation patterns of Hcm1 are fixed because all sites are changed to either phosphodeficient or phosphomimetic amino acids, the fitness of almost all mutants decreases in stress relative to WT. This raises the possibility that it is not inactivation of Hcm1 that is important for cells to maintain fitness in stress, but rather dynamic regulation conferred by phosphorylation and dephosphorylation of the TAD.

### Requirement for Cks1-dependent priming in elevated in stress

If dynamic regulation of Hcm1 is important for cells to survive in stress, we hypothesized that mechanisms that enhance phosphorylation by CDK would be more important in stress. One mechanism that promotes phosphorylation by CDK is kinase priming facilitated by the CDK accessory subunit Cks1. Cks1 binds to phosphorylated threonines at the N-terminal end of a multisite phosphorylated domain and promotes phosphorylation of CDK sites that are C-terminal to its binding site [21,22]. Importantly, Phosphosite Scanning can reveal Cks1-dependent regulation. By screening libraries that include WT phosphosites in combination with phosphodeficient or phosphomimetic mutations it is possible to identify how a mutation at one site (either an unphosphorylatable alanine or glutamic acids, which mimic the charge of a phosphate but don’t facilitate Cks1 binding) influences phosphorylation at other sites [20]. Therefore, we screened two additional libraries of Hcm1 mutants to investigate Cks1-dependent priming: one in which all phosphosites are either WT or a non-phosphorylatable alanine (WT/A library, Fig 3A); and one in which all phosphosites are either WT or phosphomimetic (WT/E library, Fig 4A) [20]. The fitness of all mutants was compared between control conditions and LiCl conditions, as described above.

**Figure 3.**
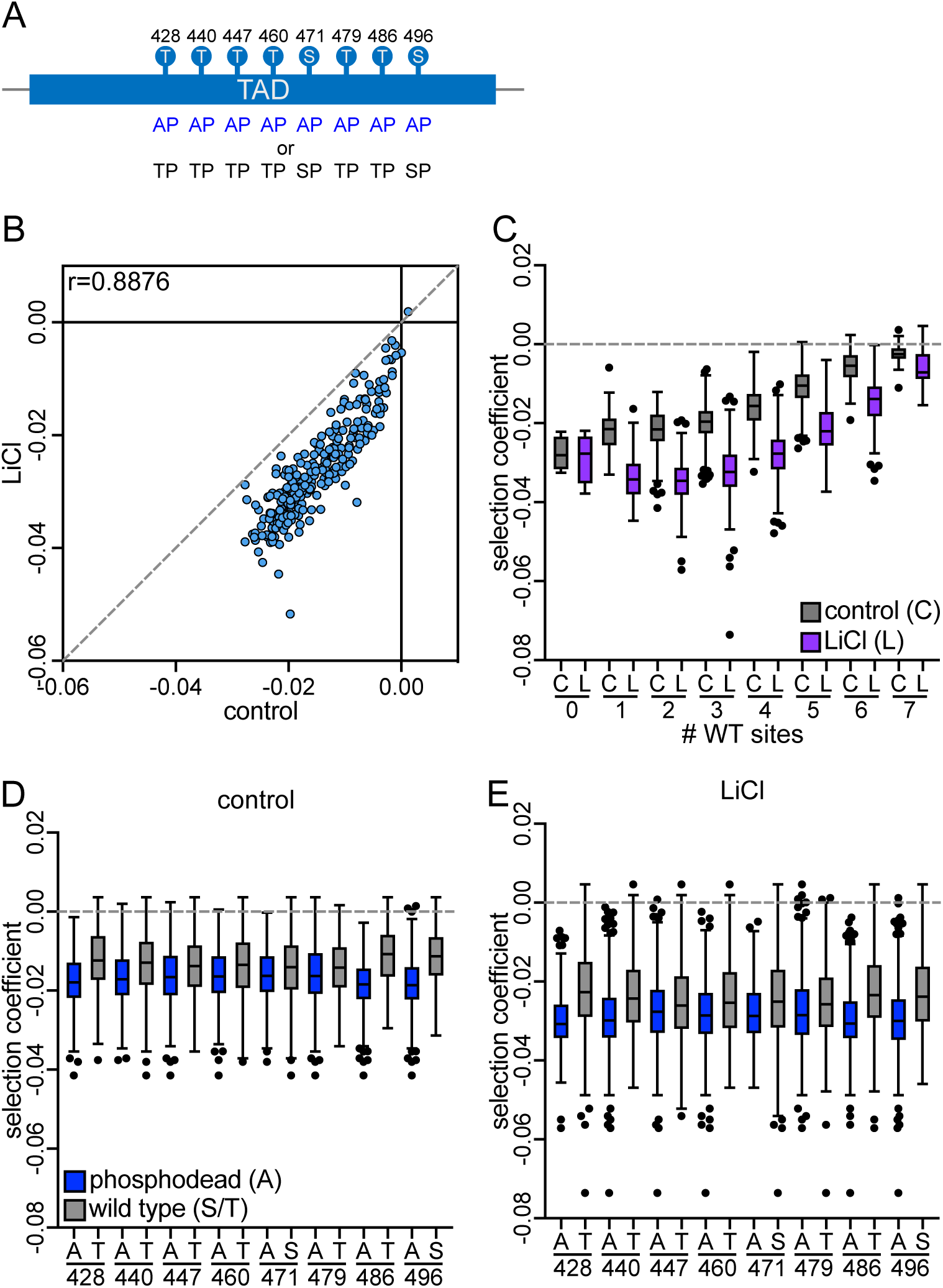
All phosphosites in the Hcm1 TAD contribute to fitness in stress. (A) Schematic depicting the Hcm1 WT/A library. (B) Scatterplot comparing average selection coefficients for each mutant in the WT/A library in control and LiCl media. Pearson correlation (r) is indicated. (C-E) Box and whisker plots comparing the selection coefficients of different groups of mutants. The black center line indicates the median selection coefficient, boxes indicate the 25^th^-75^th^ percentiles, whiskers represent 1.5 interquartile range (IQR) of the 25^th^ and 75^th^ percentile, black circles represent outliers. In all panels, selection coefficients are an average of n=4 biological replicates. (B) shows selection coefficients in cells with the indicated number of WT sites in control or LiCl conditions. (C-D) show selection coefficients of mutants that are either WT (S or T) or alanine at each position, in control (C) or LiCl containing medium (D). Statistics for panels C-E are included in Dataset S2.

**Figure 4.**
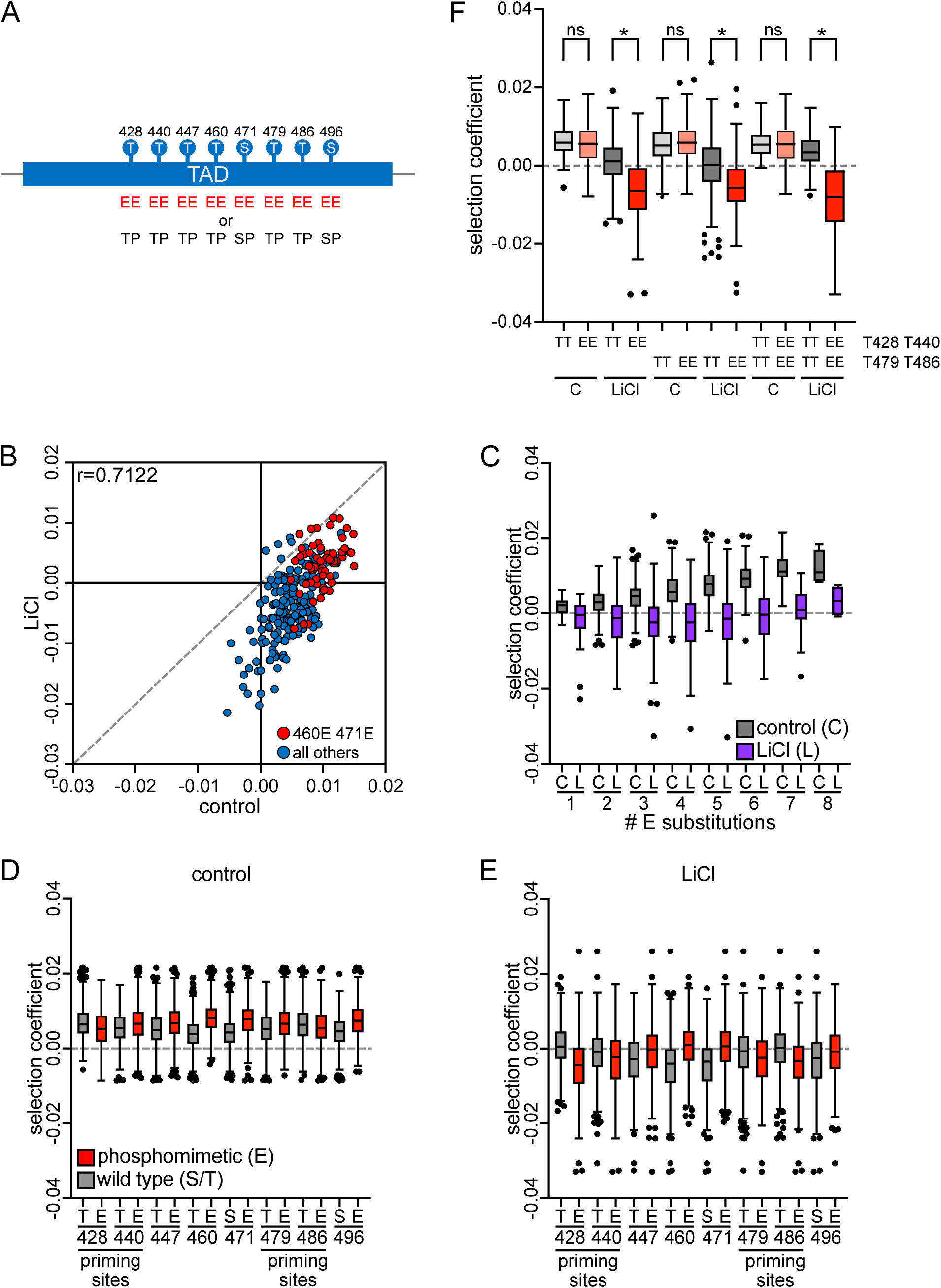
Elevated importance of processive Hcm1 phosphorylation in stress. (A) Schematic depicting the Hcm1 WT/E library. (B) Scatterplot comparing average selection coefficients for each mutant in the W/E library in control and LiCl media. Pearson correlation (r) is indicated. Red represents mutants that are phosphomimetic at sites T460 and S471, blue represents all other mutants. (C-F) Box and whisker plots comparing the selection coefficients of different groups of mutants. The black center line indicates the median selection coefficient, boxes indicate the 25^th^-75^th^ percentiles, whiskers represent 1.5 interquartile range (IQR) of the 25^th^ and 75^th^ percentile, black circles represent outliers. In all panels, selection coefficients are an average of n=4 biological replicates. (C) shows selection coefficients in cells with the indicated number of phosphomimetic mutations in control or LiCl conditions. (D-E) show selection coefficients of mutants that are either WT (S or T) or phosphomimetic at each position, in control (D) or LiCl containing medium (E). (F) shows selection coefficients for mutants that are WT (TT) or phosphomimetic (EE) at indicated positions. Statistics for panels C-F are included in Dataset S2.

First, we performed Phosphosite Scanning of the WT/A library in control and LiCl stress. Similar to the A/E screen, selection coefficients for all mutants were correlated between control and LiCl stress, however all scores were reduced in LiCl compared to control (Fig 3B). Moreover, alanine substitutions in an otherwise WT background (the W/A library) reduced fitness to a greater extent than observed when the same alanine substitutions were included an otherwise phosphomimetic background (the A/E library; compare Fig 3B with Fig 2C), which is consistent with Hcm1 regulation by Cks1 [20]. Since Cks1 can only bind to phosphothreonine [21,22], alanine substitutions reduce activity in two ways: first, activity is directly reduced because alanines cannot be phosphorylated and provide the charge conferred by a phosphate and second, activity is indirectly decreased because alanines cannot interact with Cks1 and thereby reduce phosphorylation of more C-terminal WT sites in the domain. Despite the decreased fitness of all mutants in the WT/A library, fitness generally increased with the number of WT sites in any given mutant (Fig 3C), and a WT site was more advantageous than an alanine at each position (Fig 3D-3E), in both control and LiCl conditions. These results support the conclusion that phosphorylation at each site in the TAD contributes to Hcm1 activity in both control and stress conditions.

We further explored the contribution of Cks1 priming sites to fitness in stress by screening the WT/E library (Fig 4A). Like alanine mutations, phosphomimetic (E-E) mutations cannot serve as Cks1 priming sites. Because of this, E-E substitutions effectively behave as separation of function mutations: they contribute to Hcm1 activation because they are charged, but are unable to act as Cks1 priming sites, so they impair phosphorylation of more C-terminal WT sites in the TAD. The cumulative effect of these mutations is evident when comparing selection coefficients of WT/E mutants in control and LiCl conditions (Fig 4B). Whereas most mutants in this collection are more fit than WT in control conditions, most mutants become less fit than WT in stress (Fig 4B, lower right quadrant). The subset of mutants that remained more fit than WT in stress included most mutants that have phosphomimetic mutations at T460 and S471 (Fig 4B, red circles), likely because this group of mutants does not depend on Cks1 priming to facilitate the phosphorylation of these two sites that have the greatest impact on Hcm1 activity (Fig 2E-2F). Notably, in the WT/E library, fitness did not increase with the number of phosphomimetic mutations when cells were growing in LiCl (Fig 4C), suggesting that priming by Cks1 is of increased importance when cells are growing in stress.

To further investigate the need for Cks1 priming, we analyzed the fitness of phosphomutants based on their genotype at each site. In non-stress conditions, if a phosphomimetic mutation has a negative effect and confers a reduced selection coefficient compared to the WT CDK site, it suggests that position functions as a priming site [20]. For example, in control media mutants that are phosphomimetic at site 428 are slightly less fit than mutants that are WT at the same site (Fig 4D), whereas phosphomimetic mutations at site 428 increase fitness when compared to alanine mutations in the A/E library (Fig 2E). This difference indicates that site 428 is a Cks1 priming site [20]. Notably, the deleterious fitness effect of this phosphomimetic mutation was amplified in LiCl stress, and there was a similar reduction in fitness among mutants that are phosphomimetic at site 440 (Fig 4E). Interestingly, mutants that were phosphomimetic at two additional sites, 479 and 486, also showed a reduction in fitness relative to mutants that are WT at the same sites in LiCl. Since these phosphomimetic mutations improved or had little effect on fitness in control conditions (Fig 4D), suggesting that these may be additional Cks1 priming sites within the Hcm1 TAD that only become important for fitness in stress. In fact, mutants with phosphomimetic mutations at either T428 and T440 or T479 and T468 displayed significantly reduced fitness in stress compared to mutants with WT sites, despite equivalent fitness in control conditions (Fig 4F). The increased importance of Cks1-dependent priming supports the conclusion that dynamic phosphorylation of the Hcm1 TAD is critical for fitness when cells are challenged with LiCl stress.

### Dynamic phosphorylation of the Hcm1 TAD is required for fitness in stress

If the *hcm1-8E* mutant is less fit in stress because its phosphorylation cannot be dynamically regulated, and not because it has increased activity, then an alternative mutant in which Hcm1 activity is increased by a different mechanism might not exhibit reduced fitness in stress. To test this possibility, we examined the fitness of *hcm1-3N* mutant cells. Hcm1-3N contains three alanine substitutions in the N-terminal phosphodegron that prevent proteasomal degradation and stabilize the protein [18], thereby increasing Hcm1 activity without perturbing TAD phosphorylation dynamics (Fig 5A). First, we compared the activity of the Hcm1 mutants directly, by expressing each mutant in an *hcm1Δ* reporter strain, where two copies of the Hcm1-responsive element from the *WHI5* promoter drive GFP expression. Notably, Hcm1-3N was more active than WT Hcm1, although it was less active than Hcm1-8E (Fig 5B). Despite this difference, both *hcm1-3N* and *hcm1-8E* cells exhibited similar increases in fitness in the absence of stress (Fig 5C, 1C, 5E) [15,20]. If dynamic regulation of the Hcm1 TAD is important for fitness in stress, *hcm1-3N* cells should differ from *hcm1-8E* and retain their fitness advantage.

**Figure 5.**
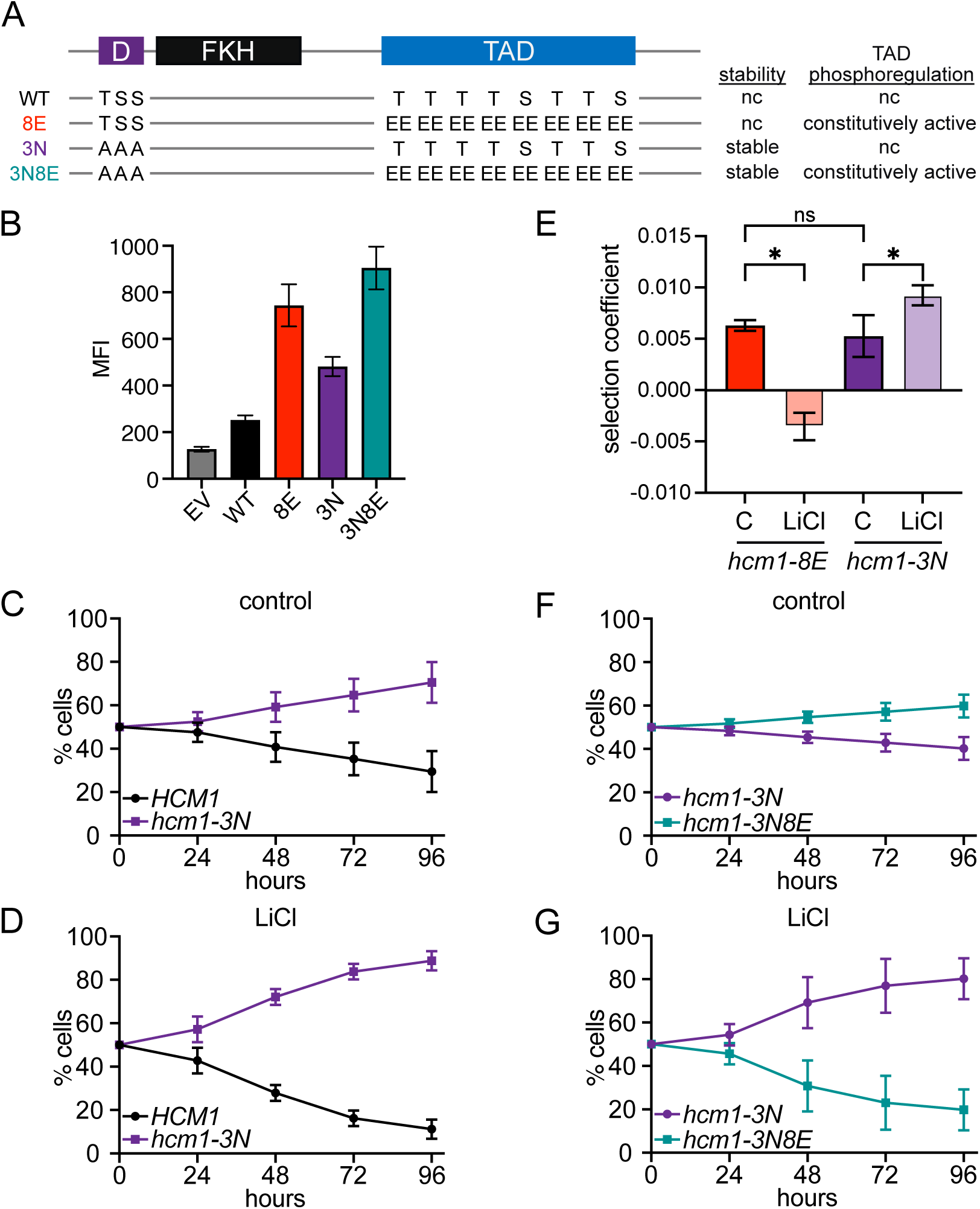
Dynamic phosphorylation of the Hcm1 TAD promotes fitness in stress. (A) Diagram of Hcm1 mutant proteins showing mutated phosphosites and impacts on protein stability and phosphoregulation of the TAD region. nc, no change. (B) Mean Fluorescence Intensity (MFI) of GFP expression in Hcm1 reporter strains expressing the indicated Hcm1 proteins or an empty vector (EV). An average of n=6 replicates is shown. Error bars represent standard deviation. (C-D) *HCM1* and *hcm1-3N* strains were co-cultured in control media (C) or media with 150mM LiCl (D). Percentage of each strain was quantified by flow cytometry at the indicated timepoints. An average of n=13 biological replicates is shown. Error bars represent standard deviations. (E) Comparison of selection coefficients of the indicated strains and growth conditions, from pairwise assays shown in Figures 1C, 1D, 4B, and 4C. One-way ordinary ANOVA with Šídák’s multiple comparisons test was used to test significance, *p<0.0001. (F-G) *hcm1-3N* and *hcm1-3N8E* strains were co-cultured in control media (F) or media with 150mM LiCl (G). Percentage of each strain was quantified by flow cytometry at the indicated timepoints. An average of n=13 biological replicates is shown. Error bars represent standard deviations.

To test this hypothesis, pairwise competition assays were carried out between *hcm1-3N* and WT cells in control and LiCl containing medium. Notably, in contrast to the decreased fitness observed in *hcm1-8E* cells challenged with LiCl stress (Fig 1D, 5E), the fitness benefit in *hcm1-3N* cells was enhanced under stress conditions (Fig 5D-5E). To compare these effects directly, these two sets of mutations were combined to generate a stable, constitutively active mutant (*hcm1-3N8E*, Fig 5A-5B). In a pairwise competition assay, *hcm1-3N8E* cells had similar fitness as *hcm1-3N* cells in control conditions (Fig 5F), as previously reported [20]. However, *hcm1-3N8E* cells were less fit than *hcm1-3N* in LiCl stress (Fig 5G). Therefore, preventing dynamic phosphorylation of the Hcm1-3N protein reverses the fitness benefit provided by its stabilization and increased expression in stress.

### Frequency modulation enhances Hcm1 activity in stress

When cells are continuously exposed to a CN-activating stress, cells experience bursts in cytosolic Ca^2+^, followed by pulses of CN activity [23,24]. This suggests that Hcm1 may undergo pulses of inactivation in chronic stress, which are then reversed by CDK activity. For some TFs, increasing the frequency of pulses of activation results in a greater induction of target gene expression, compared to increasing the amplitude of TF activity [25,26]. Therefore, we considered that Hcm1 may undergo pulses of activity in stress, through modulation of phosphorylation, which in turn could further increase the expression of its target genes. If so, WT and *hcm1-3N* cells should increase Hcm1 target gene expression in stress, compared to unstressed conditions, whereas *hcm1-8E* and *hcm1-8E3N* should not.

To test this hypothesis, we evaluated the activity of each mutant in the Hcm1 reporter strain described above, in control and LiCl stress conditions. Strains were passaged in monoculture, using the same dilution protocol that was employed for fitness assays, and GFP levels were measured using both flow cytometry and Western blotting after 40 hours of growth. As predicted, Hcm1-3N exhibited a significant increase in activity under stress, whereas Hcm1-8E and Hcm1-3N8E showed either decreased or unchanged activity (Fig 6A, S3A-S3B Fig). The WT protein also appeared more active under stress based on GFP levels detected by Western blot (S3B Fig). However, flow cytometry revealed a modest fluorescence increase in both WT and EV cells (Fig 6A, S3A Fig). Since no GFP protein was detected by Western blot in EV cells, we conclude that the fluorescence increase may be due to a slight Hcm1-independent increase in autofluorescence under stress. Notably, the increased activity of WT and Hcm1-3N could not be attributed to changes in Hcm1 protein levels (S3B Fig) or differences in the proportion of cells in S-phase (S3C Fig). These data support the model that pulses of activation increase Hcm1 target gene expression.

**Figure 6.**
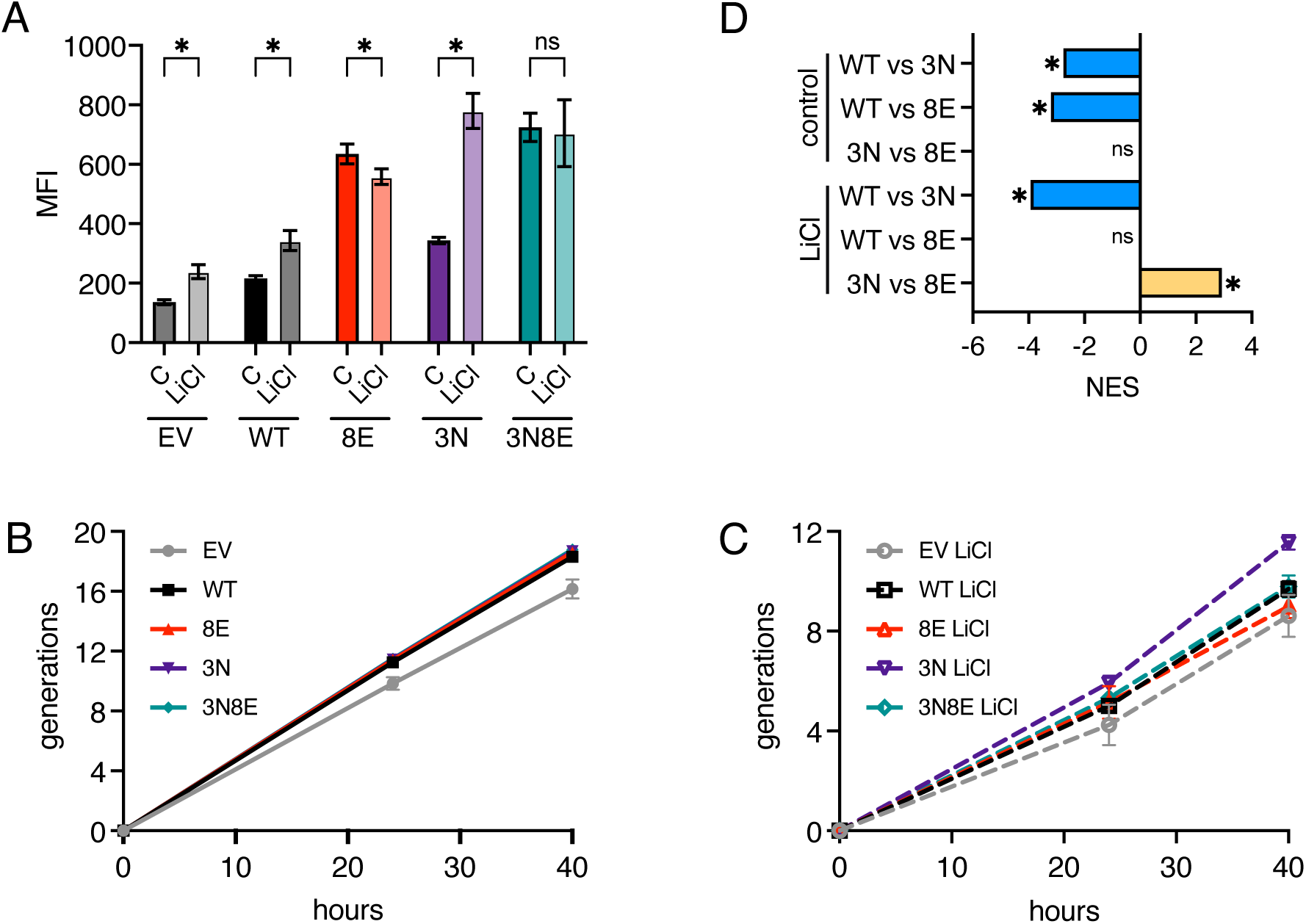
Dynamic phosphorylation increases Hcm1 activity in stress. (A) Mean Fluorescence Intensity (MFI) of GFP expression in Hcm1 reporter strains expressing the indicated Hcm1 proteins or an empty vector (EV), after 40 hours of growth in control medium or medium with 150mM LiCl. An average of n=6 replicates is shown. Error bars represent standard deviation. Two-way ANOVA with Šídák’s multiple comparisons test was used to test significance, *p<0.05. Representative histograms are shown in S3A Fig. (B-C) Cumulative generations of strains from (A) over the 40-hour experiments. Shown is an average of n=6 replicates, errors bars represent standard deviations. Strains grown in control conditions are shown in part (B). Strains grown in LiCl stress are shown in part (C). (D) Normalized enrichment scores (NES) of Hcm1 target genes from GSEA analysis of the indicated comparisons. Asterisk (*) indicates FDR=0, ns indicates not significant.

We also examined whether Hcm1 mutants could enhance proliferation by comparing population growth when the different Hcm1 proteins were expressed in the *hcm1Δ* reporter strain during the 40-hour time course. In the absence of stress, all mutants supported proliferation as well as WT Hcm1, whereas empty vector (EV) cells that do not express Hcm1 exhibited a proliferation defect (Fig 6B). In contrast, mutants differentially affected proliferation when grown in stress. Although all strains underwent fewer doublings in stress, strains expressing WT, Hcm1-8E and Hcm1-3N8E were more similar to EV cells, whereas Hcm1-3N cells displayed a proliferative advantage (Fig 6C). These results support the conclusions from pairwise competition assays (Fig 1C-1D, Fig 5C-5G) and show that Hcm1-3N expression provides a fitness advantage in stress.

We next examined how Hcm1 mutants impact the expression of endogenous Hcm1 target genes in chronically stressed cells. Hcm1 only activates target gene expression during S-phase, so the clearest way to examine the effect of Hcm1 mutants is to compare target gene expression in S-phase synchronized cells [18]. However, we found that after cells have adapted to growing in LiCl, they do not release synchronously from a G1 arrest (S4A Fig), making it difficult to quantify gene expression specifically in S-phase. Therefore, we quantified target gene expression by sequencing mRNA from asynchronous cells that were grown in control medium or LiCl for 40 hours. Because the experiment was performed in asynchronous cultures, where only ∼16% of cells are in S-phase (S4B Fig), we observed modest fold change differences between wild type and mutant strains for individual Hcm1 target genes (S3 Dataset). For this reason, we used Gene Set Enrichment Analysis (GSEA) to examine changes in expression of Hcm1 target genes as a group. GSEA revealed that Hcm1 targets were collectively expressed at elevated levels in both *hcm1-3N* and *hcm1-8E* mutants compared to WT in control conditions, confirming that the two mutants exhibit a similar increase in activity (Fig 6D, a negative normalized enrichment score (NES) indicates lower expression in WT compared to mutant). However, when cells were grown in LiCl, Hcm1 target genes increased in expression in *hcm1-3N* cells, but not in *hcm1-8E* cells. Moreover, target genes were more highly expressed in *hcm1-3N* cells than *hcm1-8E* cells when they were directly compared (Fig 6D, a positive NES indicates higher expression in *hcm1-3*N compared to *hcm1-8E*). These results demonstrate that dynamic phosphorylation of Hcm1 in stress increases expression of Hcm1 target genes and suggests that expression of these genes promotes fitness in stress.

## Discussion

Immediately upon exposure to an environmental stressor, cells rewire many cellular pathways to promote stress resistance and long-term survival. A conserved feature of this acute response is the inactivation of cell cycle-regulatory TFs and downregulation of their target genes. Although cell cycle arrest is not required for the execution of the acute stress response [27], it is possible that arrest and/or downregulation of cell cycle-regulatory genes is important for adaptation and survival after cells resume proliferation in the new environment. In support of this possibility, we found that expression of the Hcm1-8E phosphomimetic mutant, which cannot be dephosphorylated and inactivated, reduces fitness in chronic LiCl stress (Fig 1D). However, several pieces of evidence argue that simple Hcm1 inactivation does not promote fitness in stress, but rather its activity must be toggled on and off to maximally activate target gene expression and ensure fitness. First, almost all Hcm1 phosphosite mutants that retain activity but have fixed phosphorylation states are less fit, relative to WT, in LiCl than control conditions (Fig 2C). This includes mutants that have WT-like activity, as well as those that have increased activity compared to WT. Second, TP motifs that function as Cks1 priming sites to promote phosphorylation by CDK are of greater importance in stress (Fig 4F), which suggests that increased CDK-dependent phosphorylation is required to counteract CN-dependent dephosphorylation. This is seen for T428 and T440, which are previously characterized Cks1 priming sites [20], as well as two putative Csk1 priming sites T479 and T486, which are of increased importance only in stress. Finally, cells expressing the Hcm1-3N protein, which has increased activity because it is stabilized but retains phosphorylation-dependent activation [20], promotes the highest level of target gene expression and confers a fitness advantage in LiCl stress (Fig 6D, 5E). Together, these data demonstrate that Hcm1 activity is required for fitness in stress, and that the ability to add and remove phosphates is critical for maximal activity.

Dynamic regulation by phosphorylation is a recognized feature of many TFs, most notably TFs that respond to stress. In mammalian cells pulsatile nuclear localization and activation of NFAT [28], p53 [29–31] and NFκB [32–35] lead to an altered transcriptional output in response to different signals. In budding yeast, at least ten TFs display pulsatile nuclear localization in response to specific cues, thereby increasing frequency of their activation [36]. In the case of the stress-activated TF Msn2, exposure to distinct stressors triggers either sustained or pulsatile Msn2 nuclear localization [37], resulting in expression of distinct groups of target genes belonging to different promoter classes [25,26]. Notably, the CN-regulated TF Crz1 also exhibits pulsative activation via regulation of its nuclear localization [23,24]. When cells are exposed to continuous extracellular CaCl2, the frequency of cytosolic Ca^2+^ pulses increases, and these are followed by pulses of CN activation. CN then dephosphorylates Crz1, leading to pulses of nuclear localization and target gene activation. Importantly, Hcm1 does not display pulses of nuclear localization [36], instead we propose that its activation is controlled by pulses of phosphorylation. Since Hcm1 is also a CN target, when cells are growing in LiCl stress pulses of CN activity are expected to dephosphorylate and inactive Hcm1, which is then countered by CDK-dependent phosphorylation, leading to restoration of Hcm1 activity. To our knowledge the only described mechanism of TF frequency modulation is through changing TF localization, therefore dynamic phosphorylation of a TAD represents a novel mechanism of controlling TF target gene expression in stress.

Dynamic phosphoregulation is likely to also control the activities of other CDK target proteins in stress. In addition to CN, which antagonizes CDK as well as other kinases [13], the cell cycle-regulatory phosphatase Cdc14 is activated during the stress response [38–40] and could regulate phosphorylation dynamics. Moreover, a recent phosphoproteomic study monitored proteome-wide phosphorylation after acute exposure to more than 100 stress conditions and found that ∼20% of phosphorylated proteins, including Hcm1, show a change in phosphorylation after acute stress exposure [41]. Interestingly, the two Hcm1 phosphosites that changed most frequently in that study were T479 and T486, which our data suggests may be Cks1-dependent priming sites that are of increased importance in stress (Fig 5F). Notably, monitoring rapidly changing phosphorylation patterns on individual proteins is not technically feasible, therefore observations of pulsatile phosphorylation have largely been restricted to proteins where changes in phosphorylation are measured indirectly by a change in localization [36]. Here, we show that Phosphosite Scanning can be used to infer the importance of phosphorylation dynamics [20]. Screening mutants in which all sites are mutated to either non-phosphorylatable or phosphomimetic mutations reveals the importance of being able to add and remove phosphates. In addition, by screening mutants that combine phosphosite mutations and wild type sites, Phosphosite Scanning can reveal whether priming sites for the CDK accessory subunit Cks1 are important in stress, supporting the importance of dynamic phosphorylation. We anticipate that this approach will enable the investigation of phosphorylation dynamics of other proteins and reveal whether dynamic phosphorylation is a common mechanism that modulates protein function when cells are growing in stressful environments.

## Materials and Methods

### Yeast strains and plasmids

All cultures were grown in rich medium (YM-1) or synthetic media lacking uracil (C-Ura) with 2% dextrose or galactose. Cultures were grown at 30°C or 23°C, as indicated. A record of strains and plasmids used in this study can be found in S1 and S2 Tables, respectively. To construct the Hcm1 reporter strain, a synthetic DNA containing two copies of the previously characterized Hcm1 responsive element from the *WHI5* promoter [16] was cloned upstream of the minimal *GAL1* promoter and GFP. This sequence was integrated into a silent region on chromosome VI [42], and then introduced into an *hcm1Δ* background by genetic cross.

### Co-culture competition assays

Pairwise competition assays were done as described in Conti et al. 2023 ([20] and S1A Fig). Strains in which the endogenous copy of *HCM1* is regulated by a galactose inducible promoter (*GAL1p-HCM1*) and expressing either WT or non-fluorescent GFP (GFP-Y66F) were transformed with low-copy plasmids expressing WT or mutant *HCM1* from the *HCM1* promoter. Initially, cultures were grown in synthetic media lacking uracil with 2% galactose to ensure expression of *HCM1*. Logarithmic phase cells were equally mixed by adding one optical density (OD600) of each strain to the same culture tube in a final volume of 10mL C-Ura with 2% galactose. To determine the starting abundance of each strain, 0.15 optical densities were collected from the co-culture tubes. Samples were pelleted by centrifugation, resuspended in 2mL sodium citrate (50mM sodium citrate, 0.02% NaN3, pH 7.4) and stored at 4°C pending analysis by flow cytometry. To evaluate fitness in LiCl stress, co-cultures were diluted within a range of 0.005-0.04 optical densities (OD600) into synthetic media lacking uracil with 2% dextrose with or without 150mM LiCl after mixing at the start of the experiment. Cultures were then sampled and diluted every 24 hours for a total of 96 hours. Cultures reached saturation prior to dilution. At each timepoint, 0.15 optical densities were collected, pelleted, and resuspended in 2mL sodium citrate, and stored at 4°C until the conclusion of the experiment. Following the final timepoint, the percentage of GFP positive cells was quantified in each sample using a Guava EasyCyte HT flow cytometer and GuavaSoft software. 5000 cells were measured in all samples. Results were analyzed using FloJo software. Averages of n=3-13 biological replicates are shown; exact number is indicated in the figure legends. Selection coefficients for pairwise assays were calculated by calculating the slope of the best fit line of log2 fold change in mutant fraction over time, relative to WT in the same experiment. Control competitions were performed with WT (*HCM1*) cells expressing GFP or the non-fluorescent mutant GFP (GFP-Y66F). Expression of these markers had no effect on fitness (S1B-C Fig).

### Western blotting

Yeast culture amounting to one optical density (OD600) was collected, pelleted by centrifugation, and stored at −80°C prior to lysis. Cell pellets were lysed by incubation with cold TCA buffer (10mM Tris pH 8.0, 10% trichloroacetic acid, 25mM ammonium acetate, 1mM EDTA) on ice for 10 minutes. Lysates were mixed by vortexing and pelleted by centrifugation at 16,000xg for 10 minutes at 4°C. The supernatant was aspirated, and cell pellets were resuspended in 75μL resuspension solution (100mM Tris pH 11, 3% SDS). Lysates were incubated at 95°C for five minutes then allowed to cool to room temperature for five minutes. Lysates were clarified by centrifugation at 16,000xg for 30 seconds at room temperature. Supernatants were then collected, transferred to a new tube and 25μL 4X SDS-PAGE sample buffer (250mM Tris pH 6.8, 8% SDS, 40% glycerol, 20% β-mercaptoethanol) was added. The samples were incubated at 95°C for five minutes, then allowed to cool to room temperature and stored at −80°C.

For standard Western blots, resolving gels contain 10% acrylamide/bis solution 37.5:1, 0.375M Tris pH 8.8, 0.1% SDS, 0.1% ammonium persulfate (APS), 0.04% tetremethylethylenediamine (TEMED). Phos-tag gels contain 6% acrylamide/bis solution 29:1, 386mM Tris pH 8.8, 0.1% SDS, 0.2% APS, 25µM Phos-tag acrylamide (Wako), 50µM manganese chloride and 0.17% TEMED. All stacking gels contain 5% acrylamide/bis solution 37.5:1, 126mM Tris pH 6.8, 0.1% SDS, 0.1% APS and 0.1% TEMED. All SDS-PAGE gels were run in 1X running buffer (200mM glycine, 25mM Tris, 35mM SDS). Phos-tag gels were washed twice with 1X transfer buffer containing 10mM EDTA for 15 minutes (150mM glycine, 20mM Tris, 1.25mM SDS, 20% methanol) and once with 1X transfer buffer for 10 minutes on a shaking platform with gentle agitation. All gels were transferred to nitrocellulose in cold 1X transfer buffer at 0.45A for two hours. After transfer, nitrocellulose membranes were blocked in a 4% milk solution for 30 minutes.

Western blotting was performed with primary antibodies that recognize a V5 epitope tag (Invitrogen, 1:1000 dilution), PSTAIRE (P7962, Sigma, 1:10,000 dilution), G6PDH (A9521, Sigma, 1:10,000 dilution), and GFP (632381, Clontech, 1:1000). Primary antibody incubations were performed overnight at 4°C. Importantly, molecular weight makers are not shown with Phos-tag gels as they do not accurately reflect the molecular weight of proteins.

### Phosphosite Scanning screens

Phosphosite scanning screens were carried out using pooled plasmid libraries that were previously constructed and characterized [20], and transformed into a *GAL1p-HCM1* strain. A plasmid expressing WT *HCM1* was added to all libraries for normalization. During transformation, cells were cultured in YM-1 containing 2% galactose to maintain expression of endogenous *HCM1.* Following transformation, cells were cultured overnight at 23°C in synthetic media lacking uracil (C-Ura) with 2% galactose. After approximately 16 hours, an aliquot of transformed cells was removed and plated on C-Ura to confirm a transformation efficiency of at least 10X library size. Remaining cells were washed with 15mL C-Ura with 2% galactose five times, resuspended in 50mL C-Ura with 2% galactose and allowed to grow to logarithmic phase for approximately 48 hours at 30°C. The starting population was sampled to determine the initial abundance of each mutant in the population prior to selection. Cell pellets amounting to 20 optical densities were harvested, frozen on dry ice, and stored at −80°C prior to preparation of sequencing libraries. To evaluate fitness in stress, cultures were then diluted into synthetic media lacking uracil with 2% dextrose with or without 150mM LiCl after sampling at time zero. For all timepoints after time zero, cells were diluted into a range of 0.08 and 0.1 optical densities in 10mL of the appropriate media. Cultures were sampled and diluted as above every 24 hours for a total of 72 hours.

### Illumina sequencing library preps

For analysis by sequencing, plasmids were recovered from the frozen samples using a YeaStar Genomic DNA Kit (Zymo Research). Mutant *hcm1* sequence was amplified by PCR (21 cycles) using plasmid specific primers and Phusion High-Fidelity DNA polymerase (New England Biolabs). DNA fragments were purified from a 1% agarose gel using a QIAquick Gel Extraction Kit (Qiagen). Barcoded TruSeq adapters were added to the mutant fragments by PCR (7 cycles) using primers specific to the *HCM1* region fused to either the TruSeq universal adapter or to a unique TruSeq indexed adapter. Sequences of oligonucleotides that were used in library construction can be found in S3 Table. Barcoded fragments were purified from a 1% agarose gel as described above. Pooled barcoded libraries were sequenced on a HiSeq4000 platform (Novogene) to obtain paired-end 150 base pair sequencing reads. All sequencing data is available from the NCBI Sequencing Read Archive under BioProject # PRJNA1117860.

### Phosphosite scanning data analysis

Abundance of *HCM1* alleles was quantified by counting all paired-end sequencing fragments that had an exact match to an expected sequence in both reads using a custom python script. Custom scripts used to generate count tables are available on GitHub (https://github.com/radio1988/mutcount2024/tree/main/AE_type) and Zenodo (https://zenodo.org/records/13144766). Selection coefficients were calculated as the slope of the log2 fraction of reads versus time for each mutant, normalized to the log2 fraction of reads versus time of WT. All selection coefficients for all screens can be found in S1 Dataset. Box and whisker plots were generated using GraphPad Prism software. In all box and whisker plots the black center line indicates the median selection coefficient, boxes indicate the 25^th^-75^th^ percentiles, black lines represent 1.5 interquartile range (IQR) of the 25^th^ and 75^th^ percentile, black circles represent outliers. Statistical analyses for all plots of Phosphosite Scanning data are included in S2 Dataset.

### Hcm1 activity assays

Hcm1 activity was measured by expressing different Hcm1 proteins in an *hcm1Δ* reporter strain before or after exposure to stress. At the time of collection, strains were resuspended in (50mM sodium citrate, 0.02% NaN3, pH 7.4) and GFP levels in each cell were quantified using a Guava EasyCyte HT flow cytometer and GuavaSoft software. 5000 cells were measured in all samples. Mean fluorescence activity (MFI) was calculated using FloJo software. Additional aliquots of cells were saved for analysis of cell cycle position by measuring DNA content, and for analysis of Hcm1 and GFP levels by Western blot.

### RNA purification

Cells amounting to five optical densities were harvested, pelleted by centrifugation at 3000rpm for three minutes, and stored at −80°C. Cell pellets were then thawed on ice, resuspended in 400μL AE buffer (50mM sodium acetate pH 5.3, 10mM EDTA), and moved to room temperature. 40μL 10% SDS and 400μL AE equilibrated phenol was added to each sample and thoroughly mixed by vortexing for 30 seconds. Samples were heated to 65°C for eight minutes and frozen in a dry ice and ethanol bath for five minutes. Organic and aqueous layers were separated by centrifugation at max speed for eight minutes at room temperature. The aqueous layer was then transferred to a new tube. To remove any residual phenol, 500μL phenol:chloroform:isoamyl alcohol was added and thoroughly mixed by vortexing for 30 seconds. Samples were incubated at room temperature for five minutes and the aqueous and organic layers were separated by centrifugation at maximum speed for five minutes at room temperature. The aqueous layer was transferred to a new tube (∼450μL) and the nucleic acids were precipitated by adding 40μL 3M NaOAc pH 5.2 and 1mL 100% ethanol. Samples were mixed by vortexing for 15 seconds and frozen in a dry ice and ethanol bath until completely frozen. Samples were then centrifuged at maximum speed for 10 minutes at 4°C. Supernatants were decanted and pellets washed with 80% ethanol and centrifuged at maximum speed for two minutes at 4°C. Supernatants were removed, pellets allowed to dry completely and resuspended in 50μL water. DNA was degraded by treatment with DNaseI. Samples were transferred to PCR strip tubes, 10μL 10X DNaseI buffer, 2μL DNaseI and 38μL water was added to each sample, and the samples were mixed by vortexing. Samples were incubated at 30°C for 30 minutes, then cooled to 4°C in a thermocycler. 1μL 0.5M EDTA was added to each sample and mixed. Samples were then heated to 75°C for 10 minutes and cooled to 4°C in a thermocycler. Purified RNA (100μL) was then transferred to a new tube and precipitated by adding 10μL sodium acetate pH 5.2 and 250μL 100% ethanol, and frozen in a dry ice and ethanol bath until completely frozen. RNA was then pelleted by centrifugation at maximum speed for 15 minutes at 4°C. Supernatants were decanted, the pellets washed with 80% ethanol, centrifuged at maximum speed for 2 minutes at 4°C. Supernatants were decanted and pellets allowed to air dry. Purified RNA was resuspended in water. Three biological replicates were performed. Library preparation and sequencing, including polyA mRNA selection, strand specific library preparation, and paired-end 100 base pair sequencing, were performed by Innomics/BGI Americas. All sequencing data is available in NCBI GEO and is accessible through GEO accession number GSE276435.

### RNAseq analysis

RNASeq analysis was performed with OneStopRNAseq [43]. Paired-end reads were aligned to Saccharomyces_cerevisiae.R64-1-1, with 2.7.7a [44], and annotated with Saccharomyces_cerevisiae.R64-1-1.90.gtf. Aligned exon fragments with mapping quality higher than 20 were counted toward gene expression with featureCounts [45]. Differential expression (DE) analysis was performed with DESeq2 [46]. Within DE analysis, ‘ashr’ was used to create log2 Fold Change (LFC) shrinkage [47] for all possible comparisons of WT and mutant strains, in both control and LiCl conditions. Significant DE genes (DEGs) were filtered with the criteria FDR < 0.05. Gene set enrichment analysis was performed for Hcm1 targets genes using GSEA [48] on the ranked LFC. LFC analyses and GSEA results are included in S3 Dataset.

### Analysis of cell cycle position by flow cytometry

To analyze DNA content by flow cytometry, cells amounting to 0.15 optical densities were collected, fixed in 70% ethanol and stored at 4°C. Cells were then pelleted by centrifugation at 3000rpm for three minutes, resuspended in 1mL sodium citrate buffer (50mM sodium citrate, 0.02% NaN3, pH 7.4) and sonicated. Samples were then pelleted by centrifugation, resuspended in 1mL sodium citrate buffer containing 0.25mg/mL RNaseA, and incubated at 50°C for one hour. 12.5μL 10mg/mL Proteinase K was added to each tube and samples were incubated for an additional hour at 50°C. Following incubation, 1mL sodium citrate buffer containing 0.4μL Sytox green was added to each sample and samples were left at room temperature for 1 hour or 4°C overnight, protected from light, for staining. DNA content was analyzed on a Guava EasyCyte HT flow cytometer and GuavaSoft software. 5000 cells were measured in all samples. Results were analyzed using FlowJo software. Percent cell cycle progression was calculated using the following equation: % progression = ((H – 1C)/(2C-1C)) × 100, where H = histogram mean fluorescence intensity (MFI), 1C = MFI of 1C DNA content peak, and 2C = MFI of2C DNA content peak.

## Acknowledgements

The authors thank Tom Fazzio and members of the Benanti lab for insightful discussions and critical reading of the manuscript. This work was supported by National Institutes of Health grant R35GM136280 to J.A.B.

