## Supplemental Figures and Tables for "Dynamic phosphorylation of Hcm1 promotes fitness in chronic stress"

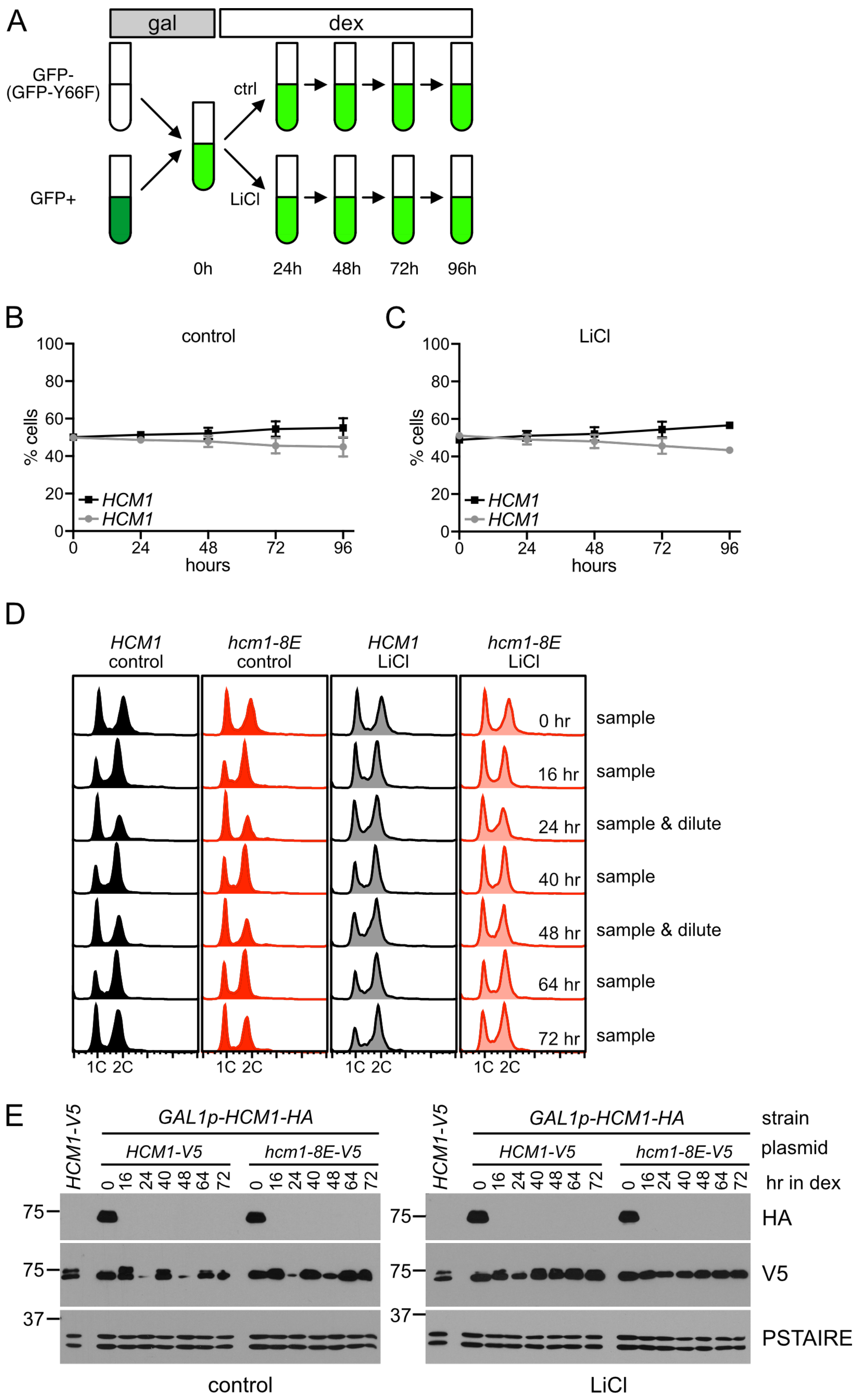

A
gal
dex
GFP-
(GFP-Y66F)
GFP+
ctrl
LiCl
0h
24h
48h
72h
96h
B
control
% cells
hours
HCM1
HCM1
C
LiCl
% cells
hours
HCM1
HCM1
D
HCM1 control
hcm1-8E control
HCM1 LiCl
hcm1-8E LiCl
0 hr sample
16 hr sample
24 hr sample & dilute
40 hr sample
48 hr sample & dilute
64 hr sample
72 hr sample
1C 2C
E
HCM1-V5
GAL1p-HCM1-HA
HCM1-V5
hcm1-8E-V5
strain
plasmid
hr in dex
HA
V5
PSTAIRE
control
LiCl

**S1 Figure. Controls for fitness experiments and Phosphosite Scanning**
(A) Schematic of pairwise competitive fitness assays. (B-C) GFP markers do not influence fitness. WT (HCM1) strains expressing GFP or GFP-Y66F were co-cultured in control medium (B) or medium containing LiCl (C). Strains have equivalent fitness. (D-E) HCM1 and hcm1-8E strains were grown in monoculture following the dilution protocol used for Phosphosite Scanning. (D) DNA content as measure by flow cytometry, which is used to infer cell cycle position. (E) Western blots showing Hcm1 expression levels. WT Hcm1 that is expressed from the genome under the control of the GAL1 promoted was detected with HA-antibodies and was undetectable after 16 hours of growth in dextrose (dex). Plasmid expressed copies of Hcm1 are tagged with V5 and were expressed throughout the experiment. In control medium, levels of Hcm1 decrease at 24 and 48 hours when cells reach saturation but increase again as cells resume cycling. Hcm1-V5 expressed from the genomic locus is included for comparison, demonstrating that the plasmid expressed copy of Hcm1 is not overexpressed. PSTAIRE is shown as a loading control.

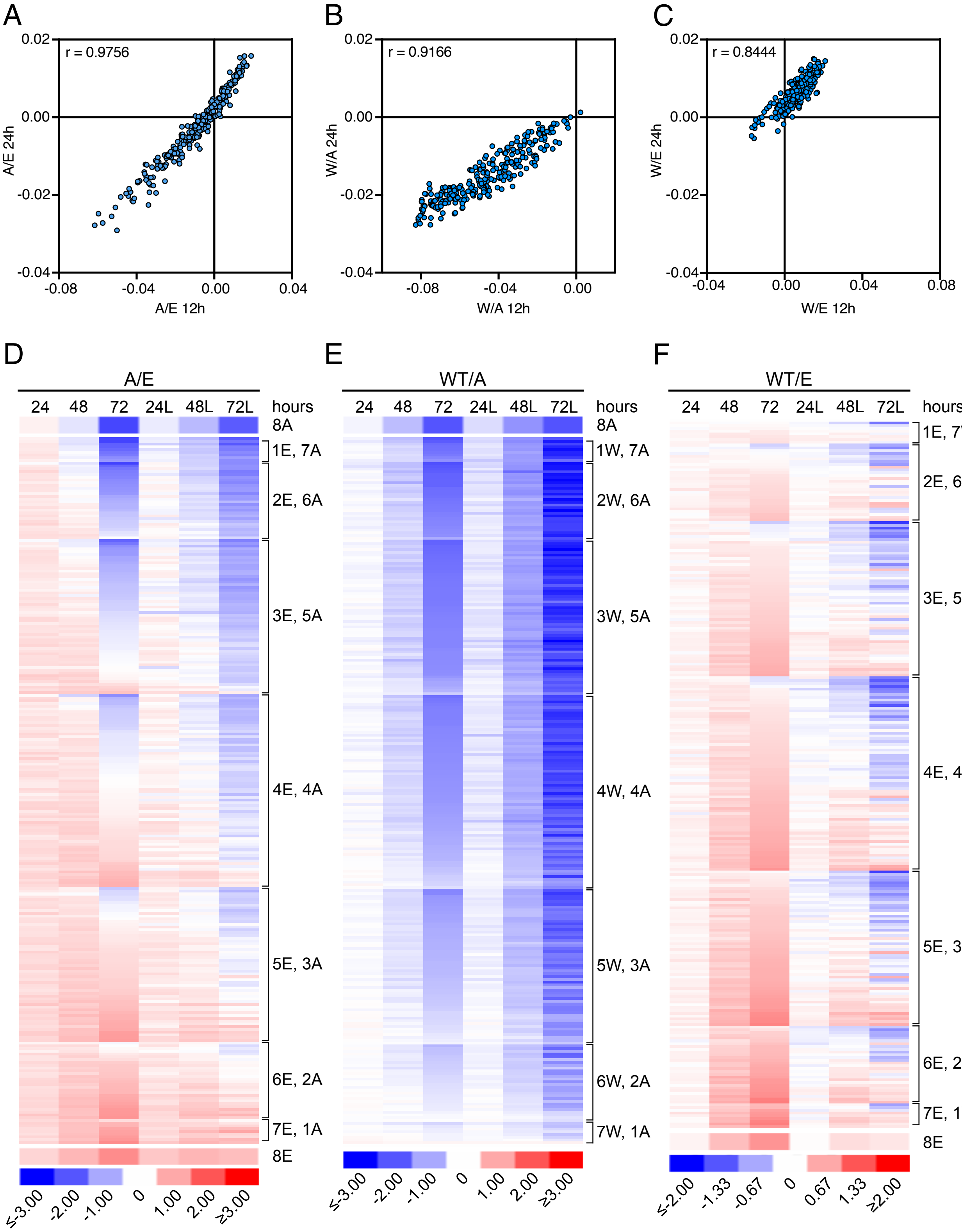
A
B
C
r = 0.9756
r = 0.9166
r = 0.8444
A/E 24h
A/E 12h
W/A 24h
W/A 12h
W/E 24h
W/E 12h
D
E
F
A/E
WT/A
WT/E
24 48 72 24L 48L 72L hours
8A
1E, 7A
2E, 6A
3E, 5A
4E, 4A
5E, 3A
6E, 2A
7E, 1A
8E
1W, 7A
2W, 6A
3W, 5A
4W, 4A
5W, 3A
6W, 2A
7W, 1A
1E, 7W
2E, 6W
3E, 5W
4E, 4W
5E, 3W
6E, 2W
7E, 1W
≤-3.00 -2.00 -1.00 0 1.00 2.00 ≥3.00
≤-2.00 -1.33 -0.67 0 0.67 1.33 ≥2.00

**S2 Figure. Supporting data for Phosphosite Scanning screens**
(A-C) Scatter plots comparing selection coefficients derived from Hcm1 Phosphosite Scanning screens carried out with dilutions every 12-hour time points, to maintain logarithmic growth, (from [20]) to those carried out with dilution every 24-hours. (A) shows screening of the Hcm1 A/E library, (B) shows screening of the Hcm1 WT/A library, (C) shows screening of the Hcm1 WT/E library. (D-F) Heat maps comparing the abundance changes of each mutant in Phosphosite Scanning screens performed in control medium or LiCl (L). Each row represents a mutant, shown is the log2 fold change in normalized read counts with respect to time zero for each mutant, all mutants have been normalized to WT. Blue indicates depletion, red indicates enrichment. Mutants are clustered by the number of phoshpomimetic mutations and ranked by increasing log2 FC within each cluster. Shown is an average of n = 4 biological replicates.

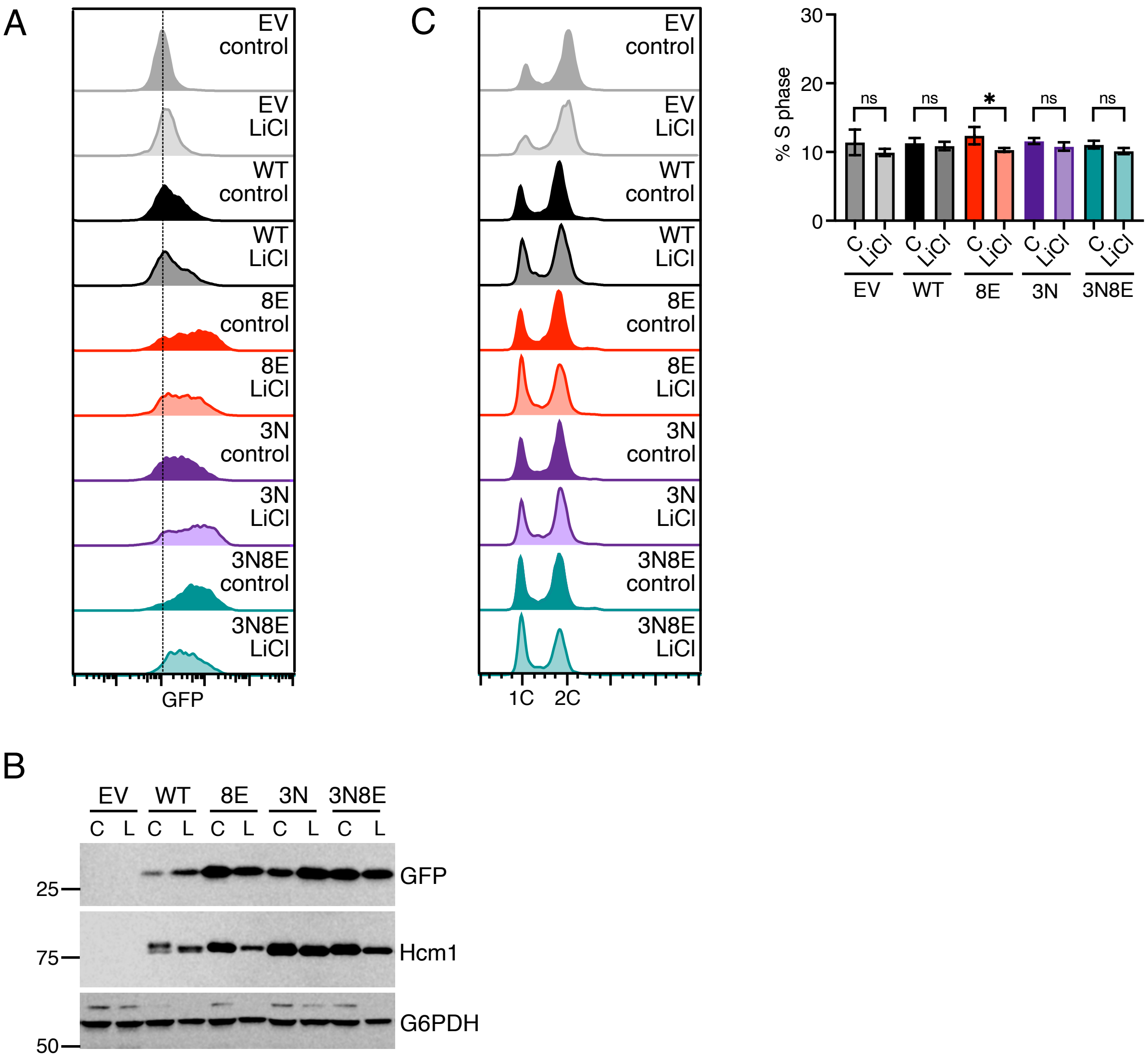


**S3 Figure. Controls for Hcm1 reporter assays**
(A) FACS plots showing the distribution of GFP expression levels in cells expressing the indicated Hcm1 mutants or an empty vector (EV), after 40 hours of growth in control medium or medium with LiCl. Shown is a representative experiment from one of n=6 replicates included in Fig 6A. (B) Western blots showing expression of GFP, Hcm1 mutants, and G6PDH (loading control) from a representative experiment from one of n=6 included in Fig 6A. (C) FACS analysis of mutants included in Fig 6A after 40 hours of growth in control of LiCl medium. Representative plots from one of n=6 experiments are shown on the left. A graph of the average percentage of cells in S-phase from all replicates is shown at the right. Error bars represent standard deviation. Significance was calculated comparing the percent S-phase in cultures in control vs LiCl medium using two-way ANOVA with Šídák's multiple comparisons test was used to test significance, *p<0.01.

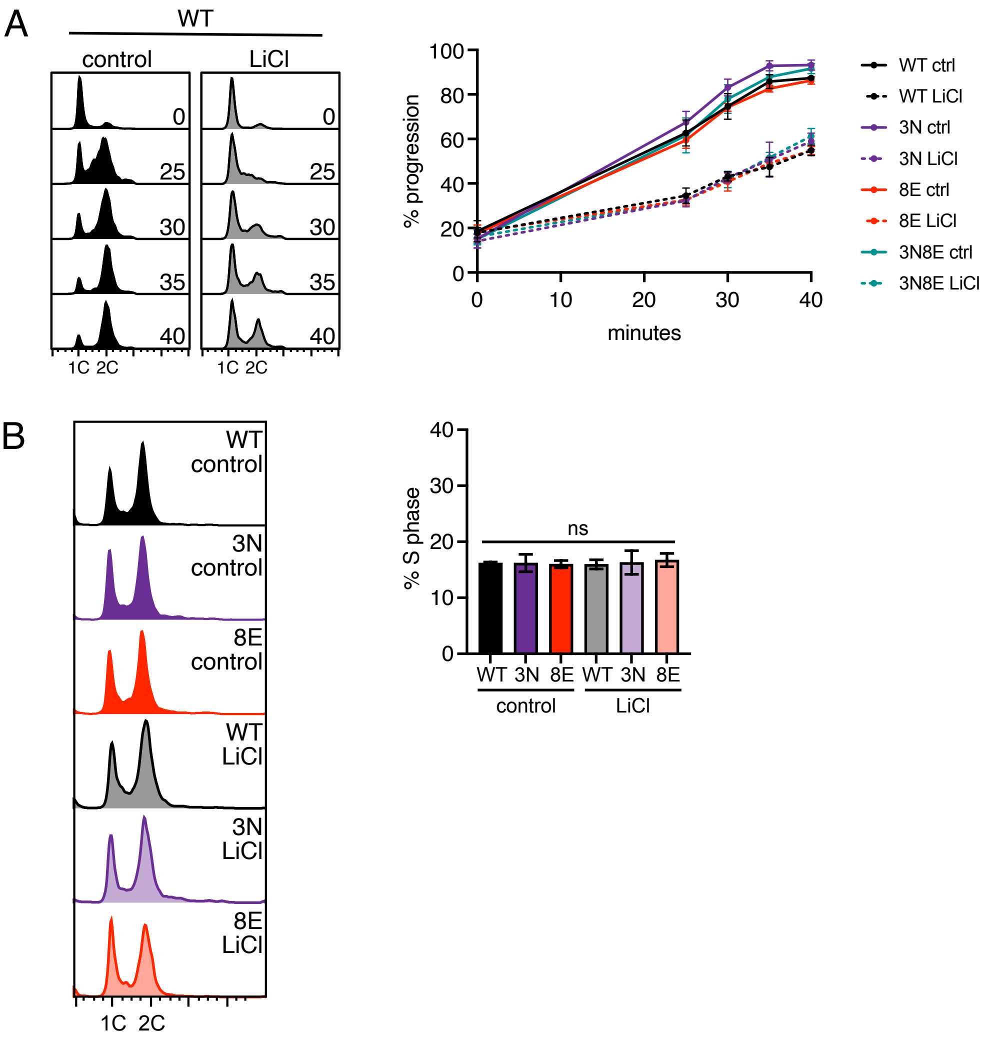


**S4 Figure. Characterization of Hcm1 mutant strains growing in stress.**
(A) WT and Hcm1 mutant cells do not release synchronously from a G1 arrest after they have adapted to LiCl stress. Monocultures of WT cells, or cells with the indicated mutants integrated into the HCM1 genomic locus, were grown in control medium or LiCl medium for 40 hours. Cells were then arrested in G1 with alpha-factor for 3 hours and then released from the alpha-factor arrest and cell cycle position was followed by flow cytometry. LiCl was kept in the medium in the indicated samples throughout the experiment. Representative plots for WT cells are shown on the left, quantitation of progression through the cell cycle is shown on the right. An average of n=3 experiments is shown, and error bars represent standard deviations. Note that all strains growing in LiCl display a similar asynchronous release and are not significantly different from one another (as determined by two-way ANOVA with Geisser-Greenhouse correction and a Dunnett's multiple comparisons test). (B) Representative FACS plots showing the DNA content of the indicated Hcm1 strains after 40 hours of growth in control or LiCl containing media (left) and percentage of S-phase cells (right). Percentage of S-phase cells is an average of n=3 biological replicates, error bars represent standard deviations. One-way ANOVA with the Geisser-Greenhouse correction and Tukey's multiple comparison test showed that none of the samples were significantly different from each other (ns).

**S1 Table. Strain table**

| Strain name | Genotype | Figure |
|---|---|---|
| YBL192 | *MATa his3∆1 ura3∆0 leu2∆0 met15∆0 HCM1-3V5-KanMX* | 1B |
| YMC50 | *MATa his3∆1 ura3∆0 leu2∆0 lys2∆0 HIS3MX6-GAL1p-HCM1-3HA-KanMX ChrVIΔ181901-182001::Hyg-TEFp-GFP + pRS316-HCM1p-HCM1-3V5* | 1C-D, 5C-G, 6D, S3, S4 |
| YMC55 | *MATa his3∆1 ura3∆0 leu2∆0 lys2∆0 HIS3MX6-GAL1p-HCM1-3HA-KanMX ChrVIΔ181901-182001::Hyg-TEFp-GFP(Y66F) + pRS316-HCM1p-hcm1-8E-3V5* | 1C-D, 5E-G |
| YMC9 | *MATa his3∆1 ura3∆0 leu2∆0 lys2∆0 HIS3MX6-GAL1p-HCM1-3HA-KanMX* | 2, 3, 4 |
| YMC53 | *MATa his3∆1 ura3∆0 leu2∆0 lys2∆0 HIS3MX6-GAL1p-HCM1-3HA-KanMX ChrVIΔ181901-182001::Hyg-TEFp-GFP(Y66F) + pRS316-HCM1p-HCM1-3V5* | 5E-G |
| YMC443 | *MATa his3∆1 ura3∆0 leu2∆0 lys2∆0 HIS3MX6-GAL1p-HCM1-3HA-KanMX ChrVIΔ181901-182001::Hyg-TEFp-GFP + pRS316-HCM1p-hcm1-3N-3V5* | 5C-E, 6D |
| YMC446 | *MATa his3∆1 ura3∆0 leu2∆0 lys2∆0 HIS3MX6-GAL1p-HCM1-3HA-KanMX ChrVIΔ181901-182001::Hyg-TEFp-GFP(Y66F) + pRS316-HCM1p-hcm1-3N-3V5* | 5C-G |
| YMC445 | *MATa his3∆1 ura3∆0 leu2∆0 lys2∆0 HIS3MX6-GAL1p-HCM1-3HA-KanMX ChrVIΔ181901-182001::Hyg-TEFp-GFP + pRS316-HCM1p-hcm1-3N8E-3V5* | 5E-G |
| YMC448 | *MATa his3∆1 ura3∆0 leu2∆0 lys2∆0 HIS3MX6-GAL1p-HCM1-3HA-KanMX ChrVIΔ181901-182001::Hyg-TEFp-GFP(Y66F) + pRS316-HCM1p-hcm1-3N8E-3V5* | 4E-F |
| YMC52 | *MATa his3∆1 ura3∆0 leu2∆0 lys2∆0 HIS3MX6-GAL1p-HCM1-3HA-KanMX ChrVIΔ181901-182001::Hyg-TEFp-GFP + pRS316- HCM1p-hcm1-8E-3V5* | 6D |
| YAR46 | *MATa ura3Δ0 his3Δ1 leu2Δ0 lysΔ0 hcm1Δ::KanMX ChrVIΔ181901-2001::HYG-Hcm2BS-GAL1p-GFP + pRS316* | 6A-C, S3 |
| YAR47 | *MATa ura3Δ0 his3Δ1 leu2Δ0 lysΔ0 hcm1Δ::KanMX ChrVIΔ181901-2001::HYG-Hcm2BS-GAL1p-GFP + pRS316- HCM1p-HCM1-3V5* | 6A-C, S3 |
| YAR49 | *MATa ura3Δ0 his3Δ1 leu2Δ0 lysΔ0 hcm1Δ::KanMX ChrVIΔ181901-2001::HYG-Hcm2BS-GAL1p-GFP + pRS316- HCM1p-hcm1-8E-3V5* | 6A-C, S3 |
| YAR50 | *MATa ura3Δ0 his3Δ1 leu2Δ0 lysΔ0 hcm1Δ::KanMX ChrVIΔ181901-2001::HYG-Hcm2BS-GAL1p-GFP + pRS316- HCM1p-hcm1-3N-3V5* | 6A-C, S3 |
| YAR51 | *MATa ura3Δ0 his3Δ1 leu2Δ0 lysΔ0 hcm1Δ::KanMX ChrVIΔ181901-2001::HYG-Hcm2BS-GAL1p-GFP + pRS316- HCM1p-hcm1-3N8E-3V5* | 6A-C, S3 |

**S2 Table. Plasmid table**

| Plasmid name | Description |
|---|---|
| pRS316-HCM1-3V5 | *HCM1p-HCM1-3V5,* CEN, URA3 |
| pRS316-hcm1-8E-3V5 | *HCM1p-hcm1-8E-3V5,* CEN, URA3 |
| pRS316-hcm1-3N-3V5 | *HCM1p-hcm1-3N-3V5,* CEN, URA3 |
| pRS316-hcm1-3N8E-3V5 | *HCM1p-hcm1-3N8E-3V5,* CEN, URA3 |
| pRS316-hcm1-A/E-3V5 | *HCM1p-hcm1-A/E-3V5, CEN, URA3* |
| pRS316-hcm1-WT/A-3V5 | *HCM1p-hcm1-WT/A, CEN, URA3* |
| pRS316-hcm1-WT/E-3V5 | *HCM1p-hcm1-WT/E, CEN, URA3* |

**S3 Table. Oligonucleotide table**

| Name | TruSeq index number | TruSeq index sequence | Orientation | Sequence |
|---|---|---|---|---|
| MC71 | Universal | N/A | FWD | AATGATACGGCGACCACCGAGATCTACACT CTTTCCCTACACGACGCTCTTCCGATCTCTC ATGGTTCGGACTTACTT |
| MC72 | 1 | ATCACG | REV | CAAGCAGAAGACGGCATACGAGATCGTGAT GTGACTGGAGTTCAGACGTGTGCTCTTCCG ATCTGGGTGCAGAGGACTTTCT |
| MC73 | 2 | CGATGT | REV | CAAGCAGAAGACGGCATACGAGATACATCG GTGACTGGAGTTCAGACGTGTGCTCTTCCG ATCTGGGTGCAGAGGACTTTCT |
| MC80 | 3 | TTAGGC | REV | CAAGCAGAAGACGGCATACGAGATGCCTAA GTGACTGGAGTTCAGACGTGTGCTCTTCCG ATCTGGGTGCAGAGGACTTTCT |
| MC81 | 4 | TGACCA | REV | CAAGCAGAAGACGGCATACGAGATTGGTCA GTGACTGGAGTTCAGACGTGTGCTCTTCCG ATCTGGGTGCAGAGGACTTTCT |
| MC82 | 5 | ACAGTG | REV | CAAGCAGAAGACGGCATACGAGATCACTGT GTGACTGGAGTTCAGACGTGTGCTCTTCCG ATCTGGGTGCAGAGGACTTTCT |
| MC83 | 6 | GCCAAT | REV | CAAGCAGAAGACGGCATACGAGATATTGGC GTGACTGGAGTTCAGACGTGTGCTCTTCCG ATCTGGGTGCAGAGGACTTTCT |
| MC84 | 7 | CAGATC | REV | CAAGCAGAAGACGGCATACGAGATGATCTG GTGACTGGAGTTCAGACGTGTGCTCTTCCG ATCTGGGTGCAGAGGACTTTCT |
| MC85 | 8 | ACTTGA | REV | CAAGCAGAAGACGGCATACGAGATTCAAGT GTGACTGGAGTTCAGACGTGTGCTCTTCCG ATCTGGGTGCAGAGGACTTTCT |
| MC86 | 9 | GATCAG | REV | CAAGCAGAAGACGGCATACGAGATCTGATC GTGACTGGAGTTCAGACGTGTGCTCTTCCG ATCTGGGTGCAGAGGACTTTCT |
| MC87 | 10 | TAGCTT | REV | CAAGCAGAAGACGGCATACGAGATAAGCTA GTGACTGGAGTTCAGACGTGTGCTCTTCCG ATCTGGGTGCAGAGGACTTTCT |
| MC88 | 11 | GGCTAC | REV | CAAGCAGAAGACGGCATACGAGATGTAGCC GTGACTGGAGTTCAGACGTGTGCTCTTCCG ATCTGGGTGCAGAGGACTTTCT |
| MC89 | 12 | CTTGTA | REV | CAAGCAGAAGACGGCATACGAGATTACAAG GTGACTGGAGTTCAGACGTGTGCTCTTCCG ATCTGGGTGCAGAGGACTTTCT |
| MC90 | 14 | AGTTCC | REV | CAAGCAGAAGACGGCATACGAGATGGAACT GTGACTGGAGTTCAGACGTGTGCTCTTCCG ATCTGGGTGCAGAGGACTTTCT |
| MC91 | 15 | ATGTCA | REV | CAAGCAGAAGACGGCATACGAGATTGACAT GTGACTGGAGTTCAGACGTGTGCTCTTCCG ATCTGGGTGCAGAGGACTTTCT |
| MC92 | 16 | CCGTCC | REV | CAAGCAGAAGACGGCATACGAGATGGACG GGTGACTGGAGTTCAGACGTGTGCTCTTCC GATCTGGGTGCAGAGGACTTTCT |

| | | | | |
|---|---|---|---|---|
| MC93 | 19 | GTGAAA | REV | CAAGCAGAAGACGGCATACGAGATTTTCAC GTGACTGGAGTTCAGACGTGTGCTCTTCCG ATCTGGGTGCAGAGGACTTTCT |
| MC94 | 20 | GTGGCC | REV | CAAGCAGAAGACGGCATACGAGATGGCCAC GTGACTGGAGTTCAGACGTGTGCTCTTCCG ATCTGGGTGCAGAGGACTTTCT |
| MC95 | 21 | GTTTCG | REV | CAAGCAGAAGACGGCATACGAGATCGAAAC GTGACTGGAGTTCAGACGTGTGCTCTTCCG ATCTGGGTGCAGAGGACTTTCT |
| MC96 | 22 | CGTACG | REV | CAAGCAGAAGACGGCATACGAGATCGTACG GTGACTGGAGTTCAGACGTGTGCTCTTCCG ATCTGGGTGCAGAGGACTTTCT |
| MC97 | 23 | GAGTGG | REV | CAAGCAGAAGACGGCATACGAGATCCACTC GTGACTGGAGTTCAGACGTGTGCTCTTCCG ATCTGGGTGCAGAGGACTTTCT |
| MC98 | 25 | ACTGAT | REV | CAAGCAGAAGACGGCATACGAGATATCAGT GTGACTGGAGTTCAGACGTGTGCTCTTCCG ATCTGGGTGCAGAGGACTTTCT |
| MC99 | 27 | ATTCCT | REV | CAAGCAGAAGACGGCATACGAGATAGGAAT GTGACTGGAGTTCAGACGTGTGCTCTTCCG ATCTGGGTGCAGAGGACTTTCT |
| MC112 | 13 | AGTCAA | REV | CAAGCAGAAGACGGCATACGAGATTTGACT GTGACTGGAGTTCAGACGTGTGCTCTTCCG ATCTGGGTGCAGAGGACTTTCT |
| MC113 | 18 | GTCCGC | REV | CAAGCAGAAGACGGCATACGAGATGCGGAC GTGACTGGAGTTCAGACGTGTGCTCTTCCG ATCTGGGTGCAGAGGACTTTCT |
| MC114 | 17 | GTAGAG | REV | CAAGCAGAAGACGGCATACGAGATCTCTAC GTGACTGGAGTTCAGACGTGTGCTCTTCCG ATCTGGGTGCAGAGGACTTTCT |
| MC155 | 28 | CAAAAG | REV | CAAGCAGAAGACGGCATACGAGATCTTTTG GTGACTGGAGTTCAGACGTGTGCTCTTCCG ATCTGGGTGCAGAGGACTTTCT |
| MC156 | 29 | CAACTA | REV | CAAGCAGAAGACGGCATACGAGATTAGTTG GTGACTGGAGTTCAGACGTGTGCTCTTCCG ATCTGGGTGCAGAGGACTTTCT |
| MC157 | 30 | CACCGG | REV | CAAGCAGAAGACGGCATACGAGATCCGGTG GTGACTGGAGTTCAGACGTGTGCTCTTCCG ATCTGGGTGCAGAGGACTTTCT |
| HCM1-PMF | N/A | N/A | FWD | CCTTCCTCTCATGGTTCGGA |
| MC70 | N/A | N/A | REV | CAAGGGATTGGGGATTCCAT |
